# Red Bull Gives You Incentive Motivation: Understanding Placebo Effects of Energy Drinks on Human Cognitive Performance

**DOI:** 10.1101/097717

**Authors:** Liane Schmidt, Pierre Chandon, Mathias Pessiglione, Hilke Plassmann

**Author notes:** **Authors contributions:** LS, PC, MP and HP designed research, LS performed research with support of the Centre Multidisciplinaire des Sciences Comportementales Sorbonne Universités-INSEAD, LS analyzed data, CP, MP and HP supervised and assisted with data analysis, LS, PC, MP and HP wrote the paper.

## Abstract

The consumption of cognitive enhancers like energy drinks (EnD) is on the rise, but do they really improve cognitive performance, and, if yes, why? We examined two novel psychological mechanisms. First, we dissociated the role of expectations and actual consumption by crossing what people consumed—Red Bull Silver Edition or a similar-tasting Sprite soda—and what they thought they consumed. We found that participants performed better in a numerical Stroop task when they believed that they had consumed an EnD, irrespective of what they had actually consume, Second, we investigated the role of motivation for such a placebo effect of EnD. We found that expected, but not actual, consumption of EnD increased the effects of incentives on cognitive performance. Our results suggest that believing that one consumes an EnD increased participants’ motivation to perform and thus enhanced their performance.

**Significance statement:** Academic doping has become increasingly popular on campuses around the globe over the last years. However, it remains unknown if and how commercially available cognitive enhancers such as energy drinks (EnD) impact cognitive performance. We varied actual and believed consumption of an EnD and measured cognitive performance and its allocation according to the magnitude of points to earn. We found that the belief that one has consumed an EnD but not its actual consumption, increased cognitive performance specifically in high point trials compared to low point trials. These findings provide novel insights into motivational processes underpinning marketing-led expectancy effects, and contribute to the current public policy debate on the use and abuse of increased alertness and concentration claims for EnD.

## Introduction

Academic doping—the consumption of stimulants, energy drinks, and other commercial products that are advertised as cognitive enhancers—has become very popular in recent years (1-4). However, research investigating the impact of prescription stimulants (4) and energy drinks (5, 6) on cognitive performance and alertness, and their psychophysiological mechanisms, has been mostly inconclusive.

The goal of this paper is to improve our understanding of whether—and if yes, how—popular commercial cognitive enhancers (i.e., energy drinks, or EnD) affect cognitive performance. Previous studies have shown that the sugar in unlabeled EnD can improve some measures of cognitive and physical performance (5, 6). However, because the drinks in these studies were administered without a label, these studies cannot disentangle to what extent mere expectations about the effects of consuming a drink labeled as EnD influence cognitive performance. This is important given the distinct taste and color of the EnD that was used in all of these studies and that thus might have revealed the experimental condition to the participants.

Interestingly, researchers have speculated that some of the effects of EnD on cognitive performance may indeed be driven by marketing-led expectancies about their effects, rather than by the effects of their actual ingredients (7, 8). Advertising for Red Bull®, for example, claims that it “improves performance and concentration, so you can tackle everything from physics to workouts.” Against this background, our first hypothesis was that the label of the drink (i.e., whether people think they consume and EnD vs. not) would increase cognitive performance.

Second, if this holds, we aimed at investigating how such positive expectancies, associated to an EnD label, would increase cognitive performance. Based on the vast interdisciplinary literature on expectancy-based placebo effects across domains (9), we focused on the role of motivation. Evidence from functional magnetic resonance imaging (10, 11), patient (12-15) and animal (16) studies showed that across domains positive expectancies linked to the administration of a sham treatment can trigger activity in dopaminergic pathways and their targets — the ventral striatum and the ventromedial prefrontal (18). These brain systems have been linked to incentive motivation (19) — a term from behavioral neuroscience that designates the ability to allocate effort according to the magnitude of incentives (20). However, the role of motivation for placebo effects has never been directly tested in past studies. Building on this convergence, we hypothesized that positive expectancies linked to thinking that one consumes an EnD (i.e., linked to an EnD label) should enhance incentive motivation and through it better cognitive performance when incentives to perform are high versus when they are low.

To test these hypotheses we dissociated the role of expectancies linked to drinking an EnD from those of its ingredients by using a balanced between-participants placebo design, which has not been done previously. In other words, we varied not only the actual drink consumed (EnD vs. placebo, a soda of similar flavor and color) but also what participants thought they consumed (EnD label vs. soda label). Participants performed a numerical Stroop task, that offered different levels of incentives to perform well, and has previously been shown to depend upon the activation of the ventral striatum (17). In addition, after training on the task and on the first trial they indicated how well they expected to perform on the task. This measure of anticipated performance allowed us to infer how participant’s expectancies varied as a function of label and drink. Taken together our design and behavioral task assessed the relative effects of label and drink on (1) cognitive performance (i.e. effort), (2) effort allocation according to the magnitude of incentives (i.e. incentive motivation), and (3) performance expectancy (i.e. anticipated performance).

## Results

### Effects of actual and believed consumption on incentive motivation

In line with our hypothesis, we found a significant main effect of EnD label on cognitive performance (ß = 0.21, SE = 0.1, *p* = .03, 95% CI [.01 – .41]) that stand in contrast to a null effect of drink (ß = −0.002, SE = 0.1, *p* = .98, 95% CI [-.2 – .19]). Importantly, the main effect of EnD label was entirely driven by an interaction of EnD label with incentives (ß = 0.05, SE = 0.02, *p* = .002, 95% CI [0.02 – 0.09]; see Table 1): As shown in Figure 1a, participants who consumed a drink labeled as an EnD showed an enhanced sensitivity to incentives. Regardless of what they had actually consumed, participants assigned to the EnD-label groups allocated their cognitive performance according to the magnitude of incentives. This enhanced incentive motivation implied that labeling the drink as an EnD facilitated cognitive performances only in trials with high incentives at stake.

**Fig. 1.**
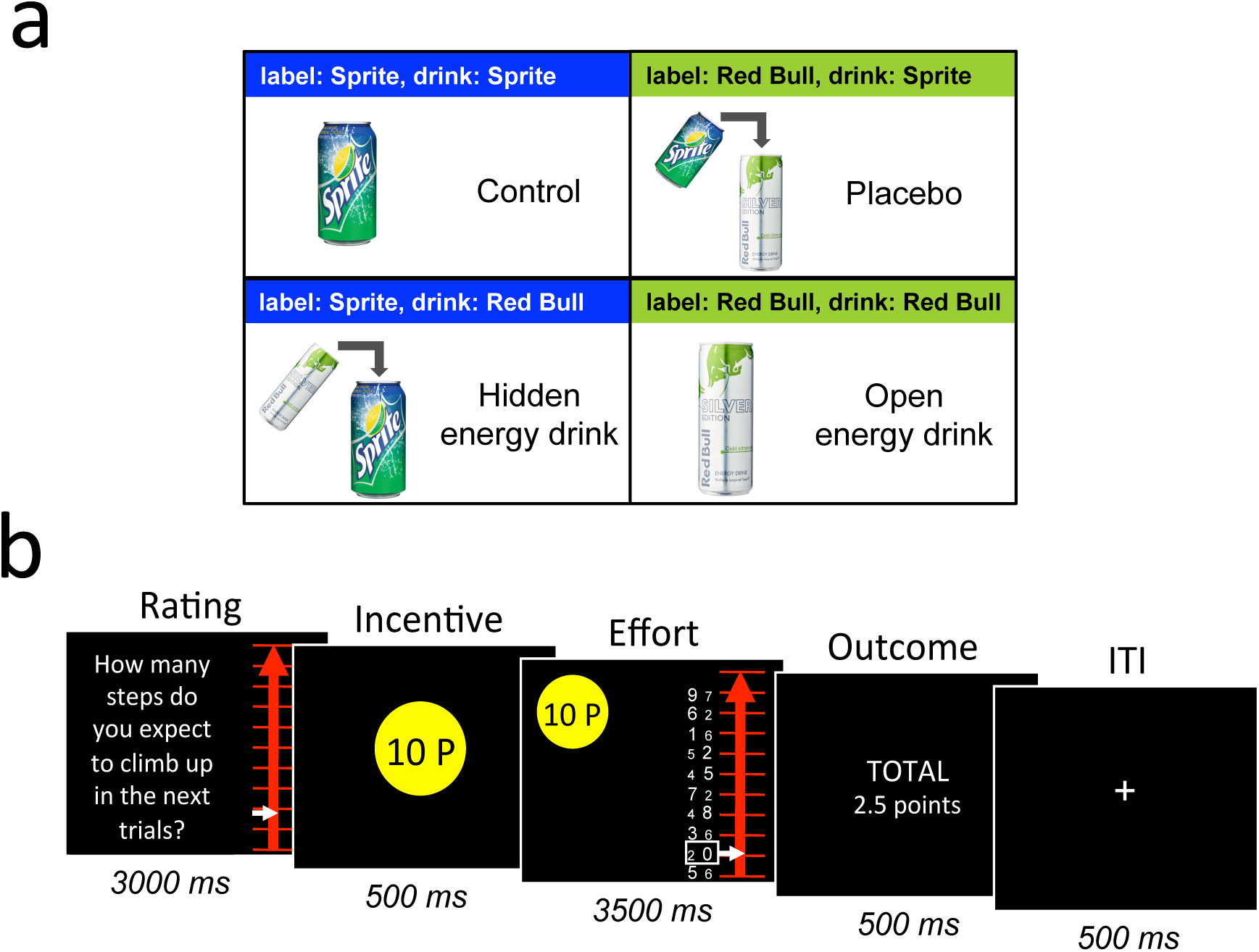
(A) Experimental design: In a 2×2 between-participant double-crossed balanced placebo design, participants were assigned to a control, a placebo, a hidden EnD, or an open EnD group. Each group counted 22 participants. (B) Task timeline: Successive screenshots displayed are shown from left to right with durations in ms. Participants rated every sixth trial how well they expected to perform during the next block of trials. The goal in each trial was to move a white cursor up the scale as high as possible in order to earn as many points as possible. Each step on the scale represented 10% of the points to earn in a given trial. To move the cursor up step by step participants had to select the keyboard button on the side of the numerically greater figure in the white box.

**Fig. 2.**
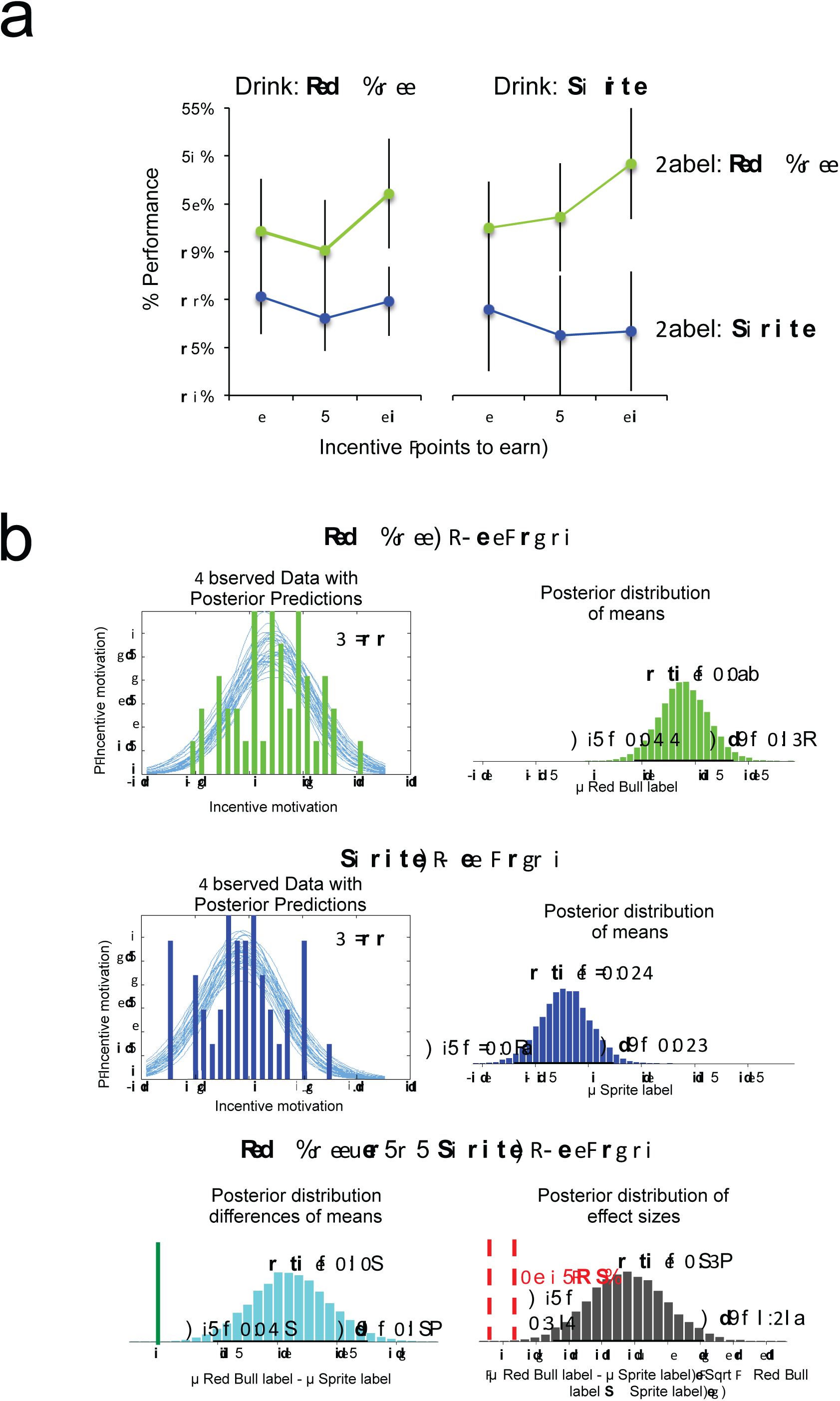
Placebo effects of Red Bull label on incentive motivation. (A) The graph depicts cognitive performance during low- (1-point), medium- (5-point), and high-incentive trials (10-point) by drink label and actual consumption condition. Error bars correspond to standard errors of the mean. (B) Bayesian estimation of the effect of EnD and soda label on incentive motivation. The upper and middle left panels depict histrograms of the observed beta estimates for the effect of incentives on cognitive performances in the EnD label group (green), Soda label group (blue) with representative examples of posterior predictive distributions superimposed. The upper and middle right panels depict the estimated posterior distributions of these effects given the observed data with mean (mode) and marginal minimal (min) and maximal (max) values. The lower panels depict the posterior distribution of mean differences and effect sizes for this mean difference. ROPE designates the region of practical equivalence (red broken bars). Note, the green bar indicates the localization of zero mean difference, which falls outside the 95% high density interval (HDI).

In addition, we applied Bayesian statistics to test the robustness of the effect of label on incentive motivation versus the null effect of drink consumed (21, 22). As shown in Figure 1b, the Bayesian estimation of the posterior distribution of credible values for the difference in mean incentive motivation between the EnD and soda label group revealed an exclusion of zero difference (mode=0.11, 95% HDI [0.047 – 0.175]). The posterior distribution of credible values for effect sizes (mode=0.735, 95% (HDI): [0.314 – 1.218]) also rejected zero (i.e. the region of practical equivalence (ROPE) counted 0% of credible mean differences). In contrast, the posterior distributions of possible differences in mean incentive motivation between the EnD and soda drink group accepted the null hypothesis of zero difference (mode=-0.001, 95% HDI: [-0.073 – 0.064]) with 34% of possible effect sizes falling within the ROPE (mode=-0.015, 95% HDI: [-0.423 – 0.435], ROPE=34%). Taken together, our results provide convergent evidence for a credible non-zero positive effect of label on motivation to perform contrasted by a possible null effect of drink.

### Effects of actual consumption and expected consumption on anticipated performance

In order to investigate whether our placebo manipulation indeed induced positive expectancies about the efficacy of the consumed drink to enhance performance, we used the very first anticipated performance rating as a measure of expectancy. A linear mixed model found a main effect of label on anticipated performance that trended toward significance (ß = 0.42, p = .06 95% CI [-0.02 – 0.86]) (SI Table 3). No fixed effect of drink and no significant interaction label by drink were observed. Bayesian estimation of the posterior distribution of possible mean differences in anticipated performance between the EnD and soda label group revealed that the 95% HDI included zero as a possible value (mode=0.079, 95 % HDI [-0.009 – 0.176]) with 6% of possible effects sizes falling within the ROPE (mode=0.372, 95% HDI [-0.042 – 0.818]). The posterior distributions for possible values for mean differences between the EnD and Soda drink group also included zero as a possible value (mode=0.028, 95% HDI [-0.062 – 0.127]). Yet, the distribution of possible non-zero effect sizes was narrower with 29% of possible values falling within the ROPE (mode=0.121, 95% HDI [-0.299 – 0.56]).

Further exploration of the data revealed that the trending difference in anticipated performance between the EnD and Soda label group was driven by a significant difference between participants, who consumed soda in a EnD can (i.e., the placebo group) and participants, who consumed soda in a soda can (i.e., the control group) (*placebo* > *control group*: *M* = 71 ± 0.4% vs. *M* = 57 ± 0.5%, t(42) = 2.03, *p* = .04 two-tailed, Cohen’s *d* = 2.8). Bayesian estimation of posterior distributions of the mean difference values between the placebo and control group revealed zero still as a possible value (mode=0.125, 95% HDI: [-0.007 – 0.272]) with a larger range of possible non-zero effect sizes involving only 5% of possible values within the ROPE (mode=0.59, 95% HDI: [-0.042 – 1.236]).

## Discussion

We investigated whether and why both actual and believed consumption of energy drinks improve cognitive performance. Our findings provide first behavioral evidence that motivational processes play a key role in placebo effects of EnD labels on performance.

Interestingly, our credible null effect of the actual drink consumed seems to be contrary to previous findings that investigated the role of EnD’s content on mental and physical performance (23, 24). Our study differs from previous work in that both the EnD and the soda contained similar amounts of sugar (Sprite: 7 g per 100 ml; Red Bull: 11 g). Our setting therefore matches what happens when people choose an energy drink rather than a soda in the hope that it will enhance their cognitive performance.

In comparison to prior work on incentive motivation for motor effort (17, 20), we observed no effects of incentives on performance in participants assigned to the soda-label groups. We cannot disentangle whether these null effects were driven by the enhanced motivation in the EnD-label groups, a decrease in motivation in the control groups, or both. Future studies measuring baseline incentive motivation prior to group assignments are needed to address this issue.

More broadly, our findings extend the previous literature on expectancy-based placebo effects in important ways. Previous studies could link dopamine functioning and reward-related behavior to placebo effects, suggesting that the activation of dopamine pathways and supposedly linked motivational processes might be the common mechanism that implements placebo effects across domains (18). Here, we provide behavioral evidence for the importance of such motivational processes for placebo effects on cognitive performance and predict that motivation might play a similarly important role for expectancy effects in other domains. We call for future neuroscientific work to causally establish the link between expectancy, motivation, and activation of dopaminergic pathways as a common mechanism underlying expectancy-based placebo effects. In turn, if motivation indeed plays an important role more generally for expectancy-based placebo effects, this could be an important insight for clinical placebo studies and, for example, shed light on new ways to improve patient well-being through motivational placebo interventions instead of mere pharmacological ones.

## Material and Methods

### Participants and experimental procedure

To test our hypotheses, we applied a 2 (drink: Red Bull Silver Edition Lime^®^ EnD or Sprite^®^ soda) x 2 (label: “Red Bull Silver Edition Lime” or “Sprite”) between-participants design: Participants were randomly assigned to one of four experimental groups (see Figure 1a): a placebo group (i.e., they drank soda labeled as EnD), an open EnD group (i.e., they drank an EnD labeled as EnD), a hidden EnD group (i.e., they drank an EnD labeled as soda), and a control group (i.e., they drank soda labeled as soda). This design allowed us to systematically disentangle the effects of expectations about consuming an EnD from the effects of actually consuming it. The placebo and open EnD groups were told that they were part of a test group for investigating the effects of an EnD on various cognitive performance tasks. The hidden EnD and control group were told that they were included in a control group for comparison to a group that had received an EnD.

The study was approved by INSEAD’s institutional review board. All participants gave written and informed consent. Eighty-eight French college students (37 males, mean age 23.6 ± 2.6 years, 22 per group) were recruited via public advertisement. They were screened for normal to corrected-to-normal vision, no history of substance abuse, any neurological or psychiatric disorder, and any medication..During the recruitment, participants were told that the study aimed to test the effects of EnDs on cognitive performance.

All participants watched the experimenter open either a can of Red Bull Silver Edition Lime (the placebo and open EnD groups) or a can of Sprite soda (the control and hidden EnD groups). The drinks were chosen based on their similarity in taste, odor, and color, verified by a blind pretest. Importantly, the administration of the placebo and hidden EnD were identical to the administration of the open EnD and control except for the following step: After opening the cans, and while out of the participants’ sight, the experimenter switched the real EnD can with another EnD can that contained soda (placebo group) or switched the soda can with another soda can containing EnD (hidden EnD group). All participants then watched the experimenter pour 250 ml of the contents into a glass and were asked to drink all of it.

A poster highlighting the benefits of the ingredients of the EnD for mental performance was displayed in the room for all four groups; the attention of the participants assigned to the placebo and open EnD groups was drawn to this poster (see SI Figure 3 for the detailed instructions, adapted from (25)). For these groups, the experimenter further highlighted that the ingredients of one can of the EnD (80 mg of caffeine, 25.5 g of sugar, and taurine) begin to stimulate the nervous system and accelerate the heart rate about 10 minutes after ingestion in order to energize mental performance. After administration, participants were accompanied to an isolated test booth to perform the various cognitive tasks. During the 10 minutes of waiting time, participants received instructions and went through several practice trials for the main task, described below.

Participants were told that they would undertake two different cognitive tasks similar to graduate exams in finance and accounting. The first task was our main task of interest; it tested whether beliefs about the drink that participants consumed affected their performance and whether these effects would be amplified by different monetary incentives at stake on a trial-by-trial basis. The second task was, in line with our cover story, similar to graduate exams in finance and accounting (see supplements for a more detailed description). Participants were compensated €12 for their participation and received an additional bonus based on their performance in the first task.

#### Numerical Stroop task

The main task consisted of a modified version of a previously used numerical Stroop task (17) that allowed us to sample (1) participants’ cognitive performance and (2) how monetary incentives affected their cognitive performance (i.e., incentive motivation) (20, 26) in a within-participant and repeated-measure design. More specifically, the task goal involved earning as much points as possible by detecting the numerically greater figure within figure pairs that differed in their physical and numerical size. Importantly, the amount of points to earn according to their performance on a given trial could vary between 1, 5, or 10 points. The points were translated into a monetary payoff at the end of the experiment. We used this task because in addition to being a reliable task to measure cognitive performance and incentive motivation, it has also been shown to engage the dopaminergic system (17, 20, 27). This is important given that our hypothesis about the role of motivation for placebo effects stems from findings about the role of the dopaminergic system to mediate placebo effects (11, 13, 14, 18).

The task was set up as follows (see Figure 1b): Participants were first asked to indicate their anticipated performance on the upcoming trials. We asked this question to measure their expectancy about the efficacy of the drink they thought they had consumed. Participants saw a graduated line representing a ladder with 10 steps, one for each figure pair, and were asked to move the cursor onto the step they expected to reach in the upcoming trials. Because participants underwent a brief training phase on this task during the waiting time for the EnD to be “active,” they were able to form expectancies about their upcoming performance. Participants also had to indicate their anticipated performance every sixth trial, for a total of 12 measures. Note that only the first response was used as a measure of the strength of participants’ expectancies about the efficacy of the drink they had consumed. The later responses were affected by their actual performance on prior trials and thus were used as an additional measure of participants’ confidence reported in the supplements.

Subsequently, the amount of points to be earned (1, 5, or 10) was displayed in a yellow circle for 500 ms. After the point display, the same ladder used during the anticipated performance rating appeared on the right side, next to the points to earn, for a response period of 3,500 ms. Each of the 10 steps of the ladder corresponded to a fraction (10%) of the points at stake. The goal in each trial was to earn as many points as possible—that is, to move the cursor as high as possible on the ladder. To climb up one step of the ladder, participants had to indicate on which side the numerically greater figure was shown by pressing the left or right keyboard button. A white box highlighted the pair of figures to be considered. If the correct button was pressed, the white cursor moved one step up, indicating that 10% of the points at stake were earned and highlighting the next figure pair to consider. If the incorrect button was pressed, the cursor could not be moved for 1 second of the response period (3,500 ms).

Importantly, the figures varied in both numerical (between 0 and 9) and physical size (between two possible fonts).^1^ In some pairs, the numerically greater figure was also physically greater (i.e., congruent pairs), whereas in other pairs it was smaller (i.e., incongruent pairs). Incongruent pairs are known to generate a Stroop effect that requires more mental effort to inhibit interference from the irrelevant information (i.e., font size). Half of the trials consisted of a 50% mix of congruent and incongruent pairs (i.e., difficult trials), whereas the other half consisted of congruent pairs only (i.e., easy trials) to vary cognitive demand (i.e., difficulty). At the end of the response window, the cumulative total of points earned was displayed to give feedback to the participants. The feedback was shown for random time intervals (3,000 ms + 100 to 1,000 ms) in order to keep participants’ attention.

In all, we applied a 2 (difficulty: 100% and 50% congruent pairs) x 3 (number of points: 1, 5, and 10) factorial within-participant task design. Each condition was repeated 12 times, randomly distributed over a series of 72 trials with a total task duration of about 10 minutes.

### Statistical analysis approach

All statistical tests were conducted with the Statistics and Machine Learning Toolbox (MATLAB 2015a, from MathWorks). A linear mixed effects model (lme) using the fitlme function in MATLAB was conducted for cognitive performance (i.e., percentage of the number of steps climbed up the ladder on each trial) as dependent variable, with fixed effects for label (coded 1 for Sprite, 2 for Red Bull), drink (coded 1 for Sprite, 2 for Red Bull), trial number (coded 1 to 72, to correct for possible learning effects), incentives (points to earn, coded 1, 2, or 3), difficulty (coded 1 for easy, 2 for difficult), and all possible two-way interactions. The model also included three uncorrelated random effects. We nested the intercept and incentives by participants (coded 1 to 88), to assess whether performance and incentive motivation varied between participants. The third random effect included the intercept nested by label (coded 1 for Sprite, 2 for Red Bull) in order to assess effects of label on performance intercept irrespective of the fixed effects. All regressors were z-scored. Results are reported in Table 1. A similar analysis was performed for mean reaction times (see SI Table 1).

In addition we used a Bayesian approach to estimate posterior distributions of parameters describing tendencies in the comparison of incentive motivation in the EnD versus Soda label; and EnD versus Soda drink groups (22) (28). This analysis was conducted using the Matlab Toolbox for Bayesian Estimation (BEST) (https://github.com/NilsWinter/matlab-bayesian-estimation/blob/master/README.md). BEST provides similar results compared to the standard linear mixed effects model, but it allows for different interpretations. Using flat priors and a Monte-Carlo-Markov-Chain (MCMC) algorithm it generates parameter values (e.g. mean differences between groups, effect sizes) from a posterior distribution of parameter values that are jointly credible given the data. Parameter values that fall within a high-density interval (HDI, 95% of the possible parameters) are considered as more credible than parameter values falling outside the HDI. Thus, this method allows consideration of the null, for example zero difference in incentive motivation between the EnD and soda drink or label groups. Specifically, we used BEST to calculate posterior distributions of two parameters: (1) the mean differences in the effect of incentives on cognitive performance between the labels (EnD vs. soda), and the drinks (EnD vs. soda), and (2) the corresponding effect sizes of these differences. To this aim we pulled together: the placebo and open EnD groups into one EnD label group; the control and hidden EnD groups into one soda label group; the open EnD and hidden EnD group into one EnD drink group; and the placebo and control group into one soda drink group. Thus, each label and drink group counted 44 participants. The effect of incentives on cognitive performance (i.e. measure for incentive motivation) corresponded to the regression coefficient (i.e. beta) of incentives obtained for each participant using a general liner model of cognitive performance predicted by incentives, cognitive demand, trial number and all possible interactions (SI paragraph 1.2. Alternative general linear model analysis, SI table 2).

## Acknowledgements

We thank Daphna Shohamy and Vasilisa Skortskova for insightful discussion, Sebastien Robin and the staff of the Centre Multidisciplinaire des Sciences Comportementales Sorbonne-Universités-INSEAD for help with data collection.

## Supplementary Material

**SI Fig. 1.**
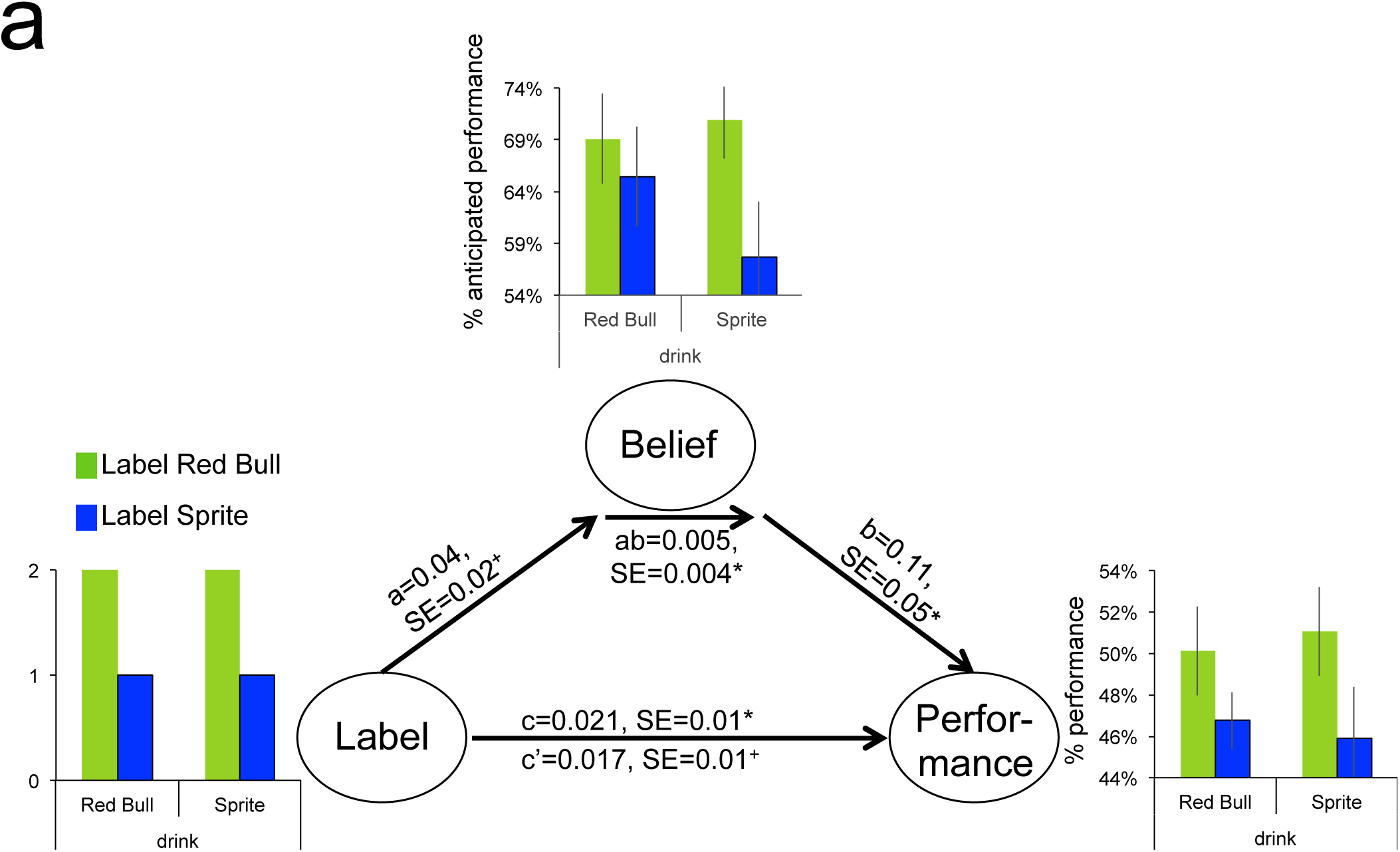
Mediation analysis diagram for the direct effect of label on cognitive performance (paths c and c'), the effect of label on expectancy (i.e., anticipated performance) (path a), the effect of expectancy on cognitive performance (path b), and the mediating effect of expectancy (path ab). * *p* < .05 two-tailed, + *p* < .05 one-tailed. Bar graphs depict effects of drink and label on belief and cognitive performance, respectively. Error bars correspond to standard errors of the mean.

**SI Fig. 2.**
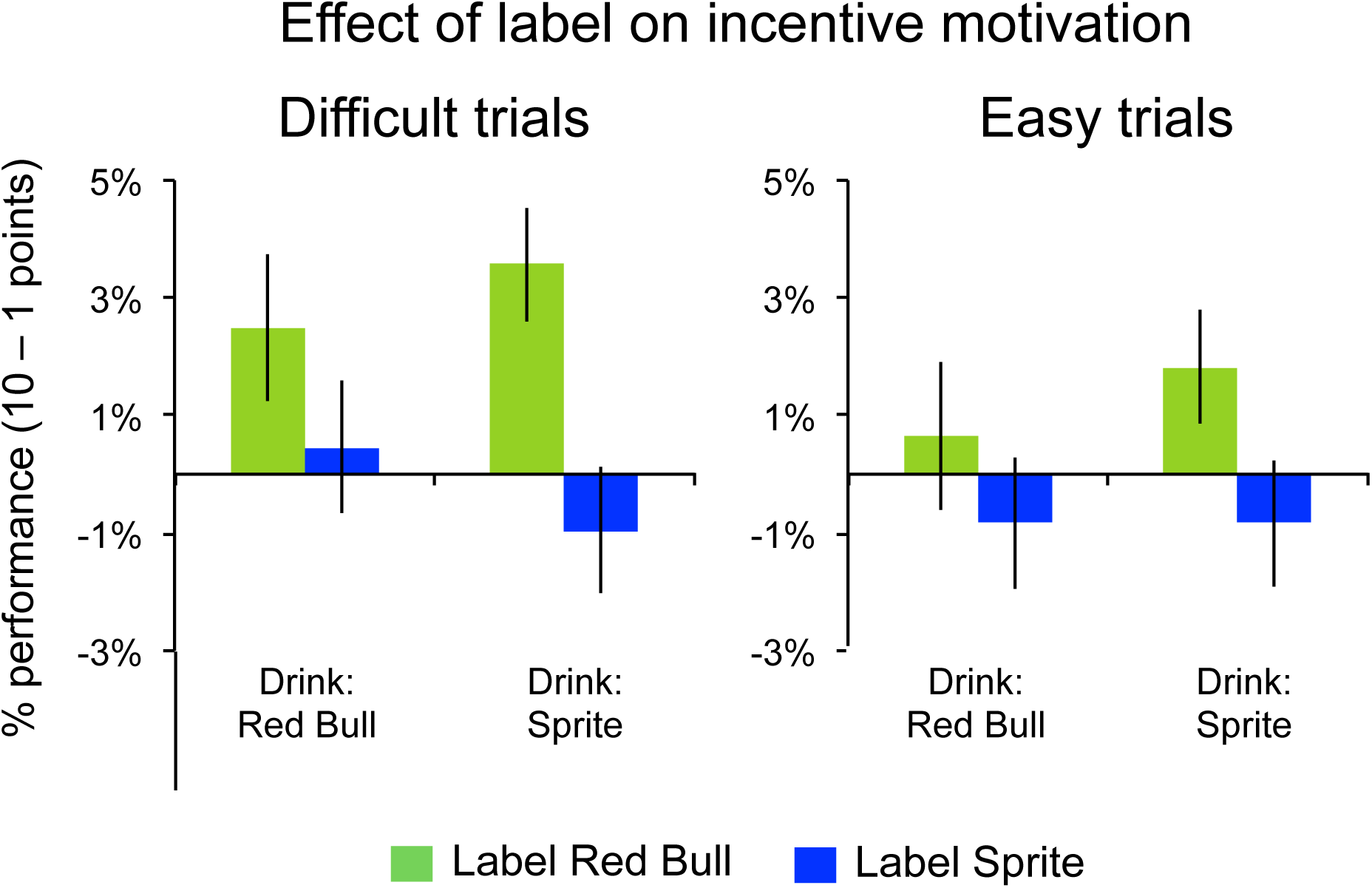
Effect of label on incentive motivation during difficult and easy trials. The bars represent the difference in cognitive performance in 10-versus 1-point trials averaged across participants in each group for difficult (left) and easy (right) trials. Error bars correspond to standard errors of the mean.

**SI Fig. 3.**
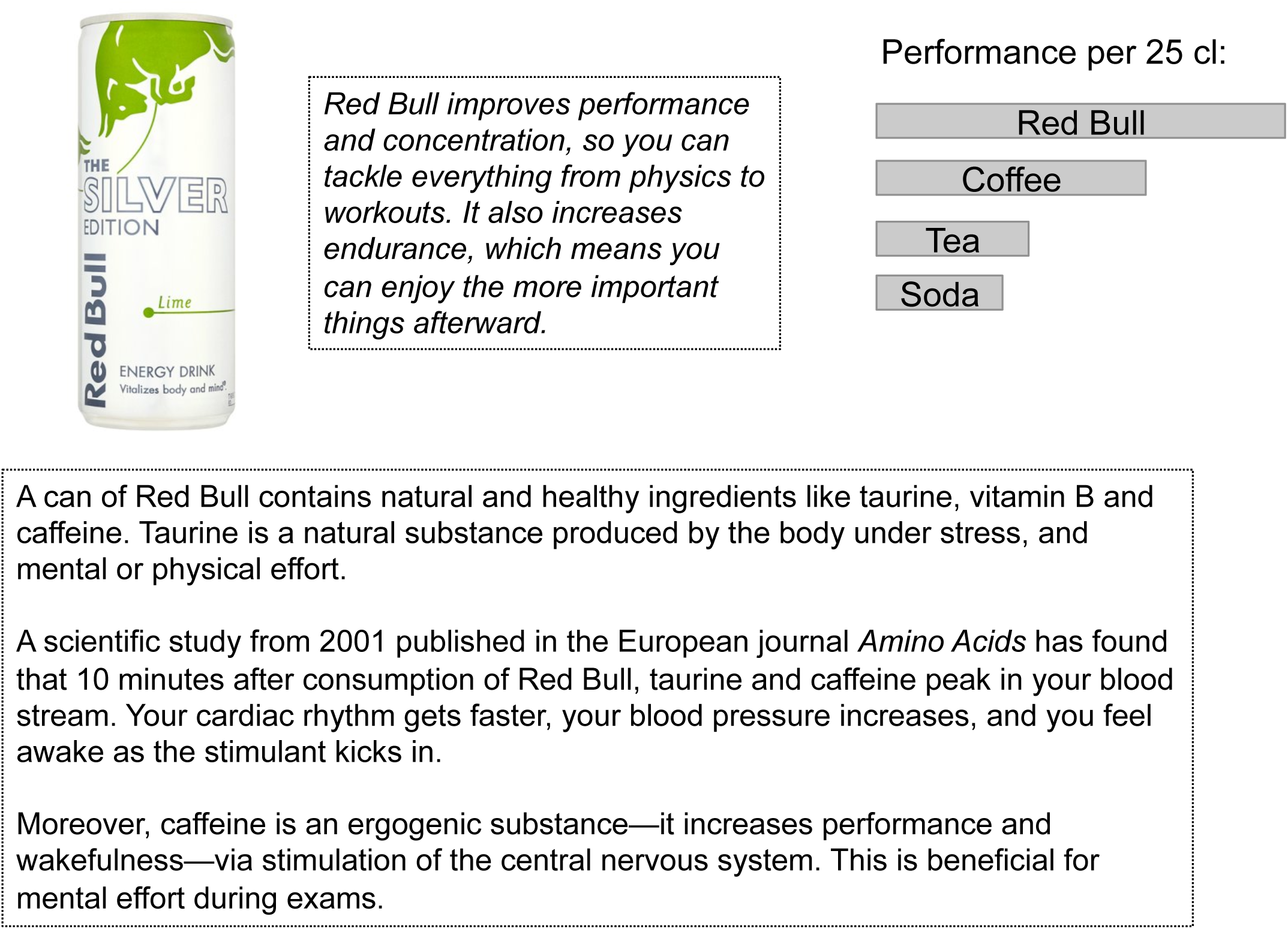
Poster highlighting the benefits of EnDs such as Red Bull for performance under pressure. Note that all participants saw this poster (in French), because it was taped on the wall of the room in which they were administered the drinks. Participants assigned to the placebo or open EnD groups were explicitly asked to read it.

## Footnotes

1 The difference in physical size (i.e., font size) was the same for all pairs, but the numerical difference varied from 1 to 5 (with two pairs of each in all trials). New number pairs were presented in each trial so that participants could not anticipate them.

